# Endothelial CD36 links arteriolar remodeling to metabolic and immune alterations in Alzheimer’s disease

**DOI:** 10.64898/2026.10.13.758947

**Authors:** Atul Kumar, Jianqing Zhang, Jada Harvey, Rachael Guenter, Rati Chkheidze, Palaniappan Sethu, Adam W. Beck, Bin Ren

**Author notes:** Correspondence to: Bin Ren, MD, PhD, FAHA, Department of Surgery, University of Alabama at Birmingham Heersink School of Medicine, Birmingham, AL 35294, USA. These authors contributed equally to this study (Atul Kumar, Jianqing Zhang, Jada Harvey).

## Abstract

Alzheimer’s disease (AD) is accompanied by profound cerebrovascular dysfunction, yet it remains unclear whether endothelial abnormalities reflect only vascular injury or altered regulatory states connecting vascular remodeling with metabolic and immune function. We integrated human brain transcriptomic and histopathological data with complementary mouse genetic and human iPSC-derived endothelial models to examine a CD36-centered endothelial-state framework in AD. Analysis of the Seattle Alzheimer’s Disease Brain Cell Atlas identified three CD36-expressing endothelial populations; intermediate-CD36 Endo 2 predominated in both reference and AD samples, although the relative subset distributions differed. In hippocampal and cortical tissue from six AD patients, endothelial CD36 expression was observed in structurally abnormal arterioles with disrupted mural organization. In complementary mouse studies, endothelial PKD1 deficiency was accompanied by increased vascular CD36 and cerebrovascular abnormalities, whereas endothelial CD36 deletion altered lipid-metabolic and innate-immune-associated transcriptional programs, including AD-relevant genes and pathways, under metabolic-inflammatory stress. To assess whether distinct human endothelial backgrounds remained pharmacologically responsive, we treated control and CRISPR-engineered PSEN1 E276A iPSC-derived ECs with GS4898, a small molecule previously linked to arterial specification. Both maintained endothelial identity but displayed different baseline transcriptional profiles and shared as well as cell-line-dependent responses to GS4898. Together, our study identifies a CD36-associated vascular phenotype in human AD, demonstrates CD36-dependent regulation of brain endothelial metabolic–immune transcriptional programs in vivo, and reveals endothelial transcriptional responsiveness to pharmacologic perturbation. The observed endothelial transcriptional plasticity may provide a basis for developing state-selective therapeutic strategies targeting cerebrovascular dysfunction in AD.

## Introduction

Alzheimer’s disease (AD) is increasingly recognized as a neurovascular disorder in which cerebrovascular dysfunction accompanies and may contribute to disease progression beyond classical amyloid-β and tau pathology. Brain endothelial cells (ECs) maintain vascular integrity, blood–brain barrier function, and neurovascular homeostasis while retaining substantial phenotypic plasticity in response to metabolic, inflammatory, and angiogenic cues.1,2 Recent single-cell and single-nucleus studies have identified disease-associated vascular states and disrupted angiogenic and barrier programs in AD, suggesting that endothelial dysfunction reflects not only structural vascular injury but also altered cellular programs.3,4 However, the mechanisms linking these endothelial changes to metabolic, immune, and vascular-remodeling abnormalities remain poorly understood.

CD36 has an important role in several of these interconnected processes. As a multifunctional scavenger receptor and fatty-acid transporter, CD36 regulates endothelial lipid uptake and metabolism, angiogenic responses, inflammatory signaling, and innate immune function.5–8 In the cerebral vasculature, CD36 has been implicated in amyloid-β-induced oxidative stress, cerebral amyloid angiopathy, and neurovascular dysfunction.9,10 ECs can also acquire context-dependent innate immune phenotypes,1,2,11,12 and metabolic reprogramming is a central feature of trained immunity and related forms of innate immune adaptation.13 These studies position CD36 as a potential link between endothelial metabolic and immune regulation in AD.

Beyond its metabolic and immune functions, CD36 also participates in endothelial vascular-state regulation. LPA/PKD1-dependent suppression of endothelial CD36 promotes angiogenic responses,8 and subsequent studies linked transcriptional repression of CD36 to proangiogenic and proarteriogenic endothelial reprogramming.14,15 Because cerebral perfusion depends on adaptive remodeling of the small-arterial network, disruption of endothelial programs that generate and stabilize functional arteriolar vessels may contribute to AD-associated cerebrovascular dysfunction. These processes can be considered within the framework of arteriolar angiogenesis (AA), a coordinated remodeling program involving endothelial state transitions, arterial/arteriolar specification, mural-cell organization, and maturation of functional small vessels.16 These observations suggest that dysregulated CD36 could couple endothelial immunometabolic dysfunction with abnormal arteriolar remodeling in AD. Furthermore, if these abnormalities reflect a regulated endothelial program rather than fixed vascular injury, they may retain the capacity for phenotypic modulation.

Accordingly, we integrated human AD single-nucleus transcriptomics and vascular histopathology with complementary human iPSC-derived endothelial and mouse genetic models to examine CD36-associated vascular, metabolic, and immune programs. We addressed three related questions: whether CD36-expressing endothelial populations and abnormal CD36-positive arterioles are observed in human AD; whether distinct human endothelial backgrounds retain transcriptional responsiveness to pharmacologic perturbation; and how endothelial PKD1 and CD36 perturbations affect cerebrovascular phenotypes and metabolic–immune transcriptional programs in vivo. To test endothelial state responsiveness, we used GS4898, a small molecule identified in a zebrafish chemical screen and previously shown to promote arterial specification and morphogenesis.17,18

## Methods

### Study design and ethics

The UAB Institutional Review Board determined that the use of de-identified human brain tissue specimens did not constitute human subjects research (IRB-300014983). Animal procedures were performed in accordance with the National Institutes of Health Guide for the Care and Use of Laboratory Animals and were approved by the UAB Institutional Animal Care and Use Committee. Publicly available, de-identified single-nucleus RNA-sequencing data were analyzed as described below.

### Human Alzheimer’s disease brain tissue and vascular histology

Formalin-fixed, paraffin-embedded hippocampal tissues from three AD patients and cortical tissues from three additional AD patients were examined for vascular morphology and endothelial CD36 expression. Sections were deparaffinized, rehydrated, and subjected to heat-induced antigen retrieval. For chromogenic immunohistochemistry, endogenous peroxidase activity and nonspecific binding were blocked before incubation with primary antibodies, followed by horseradish peroxidase-based detection with 3,3′-diaminobenzidine and hematoxylin counterstaining. For immunofluorescence, sections were blocked and incubated with primary antibodies, including anti-CD36 (clone 63; Millipore, MAB1258) and anti-α-smooth muscle actin (clone 1A4; Sigma-Aldrich, A2547), and anti-ACKR1/DARC (rabbit polyclonal; Thermo Fisher Scientific, PA5-112940),followed by appropriate fluorophore-conjugated secondary antibodies. CD36 distribution and vascular smooth-muscle organization were evaluated in cerebral arterioles, including α-smooth muscle actin (α-SMA) staining and nuclear counterstaining with DAPI where indicated. Brightfield and fluorescence images were acquired under matched settings within each experiment.

### SEA-AD single-nucleus RNA-sequencing analysis

Publicly available single-nucleus RNA-sequencing data from the Seattle Alzheimer’s Disease Brain Cell Atlas (SEA-AD) were used to evaluate endothelial CD36 expression in human AD. Integrated 10x Genomics datasets from the middle temporal gyrus, including reference/control and AD samples, were accessed through the Allen Institute Brain Map/SEA-AD resource. Endothelial cells were examined for CD36 expression, and CD36-positive endothelial populations were evaluated across the Endo 1, Endo 2, and Endo 3 subsets represented in the dataset. The relative contributions of these endothelial subsets were compared descriptively between reference and AD groups. Available subject-level metadata, including APOE4 status, Braak stage, and dementia status, were visualized across the endothelial embedding for exploratory interpretation.

### Mouse models and brain histology

Two endothelial genetic models were used. Endothelial cell (EC)-specific PKD1-deficient mice were examined for cerebrovascular morphology and endothelial CD36 expression. Brain tissues were fixed, paraffin embedded, sectioned, and evaluated by routine histology and immunostaining to assess vascular architecture, collagen deposition, and endothelial CD36 expression. For vascular immunofluorescence, antibodies included anti-CD36 (clone 63; Millipore, MAB1258) and anti-VEGFR-2 (rabbit polyclonal; Sigma-Aldrich, SAB4501645). Appropriate littermate or genotype controls were analyzed in parallel. In separate experiments, EC-specific CD36 knockout translating ribosome affinity purification (TRAP) mice and TRAP control mice were subjected to a high-fat diet for seven weeks before collection of brain tissue for endothelial transcriptomic analysis. The high-fat diet was used as a defined metabolic-inflammatory challenge to expose CD36-dependent endothelial programs and was not intended to reproduce the full AD milieu. Mice were maintained under standard housing conditions with free access to food and water.

### Endothelial TRAP and isolation of ribosome-associated RNA

Ribosome-associated mRNA was isolated directly from brain endothelial cells using a TRAP-based procedure. Brains were rapidly dissected into ice-cold buffers containing HBSS, 2.5 mM HEPES-KOH (pH 7.3), 35 mM glucose, and 4 mM NaHCO3. Tissue was transferred to prechilled extraction buffer containing 10 mM HEPES (pH 7.4), 150 mM KCl, 5 mM MgCl2, 0.5 mM dithiothreitol, 100 μg/mL cycloheximide, protease inhibitors, and RNase inhibitors and homogenized on ice. Lysates were centrifuged at 2,000 × g for 10 min at 4 °C. NP-40 and DHPC were added to the supernatant to final concentrations of 1% and 30 mM, respectively, followed by incubation on ice and centrifugation at 20,000 × g for 10 min at 4 °C. GFP-tagged ribosome complexes were recovered by immunoprecipitation with anti-GFP antibody-coupled magnetic beads using the established TRAP workflow. Beads were washed five times with high-salt wash buffer containing 10 mM HEPES (pH 7.4), 350 mM KCl, 5 mM MgCl2, 1% NP-40, 0.5 mM dithiothreitol, and 100 μg/mL cycloheximide. RNA was released in lysis buffer containing β-mercaptoethanol and purified for sequencing. The TRAP workflow was based on established endothelial and lineage-specific ribosome-affinity purification methods.19,20

### RNA sequencing and transcriptomic pathway analysis

RNA sequencing was performed by the UAB Heflin Center for Genomic Sciences. Raw FASTQ files were adapter-trimmed with Trimmomatic, aligned to the reference genome with HISAT2, and processed with SAMtools. Differential gene expression was analyzed with DESeq, and RPKM-normalized expression values were used for downstream visualization and pathway-level analyses. Genes showing at least a twofold change relative to TRAP control samples were carried forward for focused functional analyses. Gene Ontology and Kyoto Encyclopedia of Genes and Genomes pathway analyses were performed with DAVID. Gene set enrichment analysis was used to evaluate coordinated inflammatory, cytokine, innate-immune, and metabolic programs using 1,000 permutations; normalized enrichment score, nominal P value, and false-discovery rate (FDR) were used for interpretation, with FDR ≤ 0.25 and nominal P < 0.05 used for the current pathway analyses. Trained-immunity-associated genes were additionally evaluated with the trained-immunity database (TIDB).21 Enrichr-KG was used to explore functional relationships among selected interleukin and Toll-like receptor signaling components, and gene-interaction networks were visualized with Cytoscape and the Network Data Exchange platform.

### iPSC lines and maintenance

The control and PSEN1 E276A iPSC lines were provided by the laboratory of Dr. Palaniappan Sethu. The control line was originally obtained from Cellular Dynamics International and was derived from cardiac fibroblasts (CMC-100-010-001). The PSEN1 E276A line was obtained from The Jackson Laboratory (PSEN1 E276A SNV/WT; JIPSC001226). It is a heterozygous CRISPR/Cas9-engineered derivative of the KOLF2.1J human iPSC line (PSEN1 E276A/+). The control and PSEN1 E276A lines have distinct genetic backgrounds and are not isogenic. Cells were maintained on Geltrex-coated surfaces in mTeSR Plus medium and were exposed to 10 μM Y-27632 during initial plating and recovery. Medium was changed approximately every 48 h, cells were passaged near 90% confluence, and endothelial differentiation was initiated when cultures reached approximately 90–100% confluence.

### Differentiation of iPSCs into endothelial cells

Endothelial differentiation was performed using a staged protocol. Two days before induction (day ™2), iPSCs were dissociated and seeded onto Geltrex-coated six-well plates in mTeSR medium containing 10 μM Y-27632. On day 0, endothelial differentiation was initiated by washing the cells with DPBS and culturing them for 24 h in DMEM/F12 containing B27 supplement without insulin and 10 μM CHIR99021. On day 1, medium was replaced with DMEM/F12 containing B27 without insulin, 100 ng/mL VEGF, 10 ng/mL TGF-β1, and 25 ng/mL erythropoietin for 48 h. On day 3, cells were harvested and transferred to fibronectin-coated culture flasks in EGM2-MV medium supplemented with B27, 100 ng/mL VEGF, and 10 μM SB-431542 for 24 h. On day 4, medium was replaced with EGM2-MV containing B27, 200 ng/mL VEGF, and 15 μM SB-431542 for an additional 3 days. The expanded iPSC-derived endothelial cells (iPSC-ECs) were then maintained on fibronectin-coated surfaces in EGM2-MV medium supplemented with B27, 100 ng/mL VEGF, and 10 μM SB-431542.

### Immunofluorescence validation of endothelial identity

For immunofluorescence validation of endothelial identity, iPSC-ECs were cultured in chamber slides, fixed with 4% paraformaldehyde (Electron Microscopy Sciences, 15710), and blocked with 5% bovine serum albumin in PBS containing 0.1% Triton X-100. Cells were stained with Alexa Fluor 488-conjugated anti-CD31 (clone 89C2; Cell Signaling Technology, 4277S). Nuclei were counterstained as indicated. Images were acquired with a BZ-X series imaging system (Keyence) using matched acquisition settings within each experiment. CD31 staining and endothelial morphology were used as phenotypic validation of the differentiated cultures.

### GS4898 treatment of iPSC-derived endothelial cells

GS4898 was previously characterized in studies of arterial specification and morphogenesis.17,18 Control and PSEN1 E276A iPSC-derived endothelial cells (iPSC-ECs) were treated with vehicle (DMSO), 2 μM GS4898, or 5 μM GS4898 for 72 h under the maintenance conditions described above. Three independent differentiation experiments from the same control and PSEN1 E276A iPSC lines were used for the current transcriptomic and qPCR analyses. Thus, differentiation set was treated as a biological experimental block rather than as an independent donor.

### Immunoblotting and densitometric analysis

For acute kinase-signaling analysis, control and PSEN1 E276A iPSC-derived endothelial cells were exposed to GS4898 for 30 min under the maintenance conditions described above. Cells were lysed in RIPA buffer (Thermo Fisher Scientific) supplemented with protease and phosphatase inhibitors. Proteins were separated using Mini-PROTEAN TGX precast gels and transferred with Trans-Blot Turbo transfer packs (Bio-Rad). Membranes were incubated overnight at 4 °C with antibodies against total Akt (Cell Signaling Technology, 9272), phospho-Akt Thr308 (9275), phospho-Akt Ser473 (9171), total ERK1/2 (9102), and phospho-ERK1/2 Thr202/Tyr204 (4370), followed by appropriate secondary antibodies for 1 h after washing with PBST. Signals were detected with ECL substrate (Cytiva/Amersham, RPN2236) and imaged with a ChemiDoc MP system (Bio-Rad). Band intensities were quantified by densitometry using Quantity One software (Bio-Rad). Phosphorylated kinase signals were evaluated relative to the corresponding total-protein signal, and total-protein abundance was analyzed separately where indicated. Repeated immunoblot runs from the same lysate were treated as technical rather than independent biological replicates.

### RNA isolation and RT-qPCR

Cultured iPSC-ECs were washed with PBS and lysed directly in six-well plates with TRIzol reagent. Total RNA was isolated using the Direct-zol RNA Miniprep Plus kit (Zymo Research) and further purified and concentrated with the RNA Clean & Concentrator system (Zymo Research). RNA concentration was measured with a DS-11 NanoSpectrophotometer (DeNovix). For cDNA synthesis, 125 ng total RNA was reverse-transcribed using the SuperScript III First-Strand Synthesis System (Invitrogen). Quantitative PCR was performed using SYBR Green ROX qPCR Master Mix. Gene expression was normalized to GAPDH and expressed relative to the matched vehicle condition. Technical replicates were averaged within each differentiation experiment before biological-level analysis.

### NanoString nCounter gene-expression profiling and analysis

For NanoString profiling, 150 ng total RNA per sample was hybridized to the Human Tumor Signaling 360 Panel (NanoString Technologies/Bruker Spatial Biology). The current analysis included 18 samples: control and PSEN1 E276A iPSC-ECs treated with vehicle, 2 μM GS4898, or 5 μM GS4898 across three independent differentiation experiments. nCounter data processing and quality control followed the established laboratory workflow. Principal component analysis was used to examine global transcriptional structure. For the three-set reanalysis used in the present results, normalized gene-expression values were analyzed with a blocked linear model that included biological differentiation set, cell-line background, GS4898 dose, and the cell-line-background × dose interaction. Benjamini-Hochberg correction was applied across the 734 genes tested within each transcriptomic contrast. Cell-line-background × treatment interaction terms were used to identify genes showing differential responses to GS4898 between the two iPSC-EC lines. For the supplementary endothelial-identity analysis, vehicle-treated samples were additionally examined for canonical endothelial markers available on the panel (including CDH5, VWF, KDR, FLT1, TEK, NOS3, and CD34) and for brain/barrier-associated vascular genes (including TJP1, SLC2A1, ABCG2, FZD4, and EPAS1).

### Image analysis

Microscopy images within a given experiment were acquired using consistent exposure and acquisition parameters. Quantification of staining intensity, positive area, endothelial signal, or smooth-muscle coverage was performed with ImageJ/Fiji as appropriate. Representative images were selected from independent biological samples and were not used as substitutes for quantitative analysis.

### Statistical analysis

Statistical analyses for non-transcriptomic experiments were performed with GraphPad Prism version 9 unless otherwise specified. Technical replicates were averaged within each biological sample or differentiation experiment before statistical testing. Data presentation and the statistical test used for each experiment are specified in the corresponding figure legends. A two-sided P value < 0.05 was considered statistically significant for focused comparisons. NanoString transcriptome-wide analyses used Benjamini–Hochberg false-discovery-rate correction as described above.

## Results

### CD36-expressing endothelial heterogeneity and abnormal arteriolar morphology in AD

To define the distribution of CD36-expressing endothelial populations in human AD, we analyzed single-nucleus RNA-sequencing data from the Seattle Alzheimer’s Disease Brain Cell Atlas using middle temporal gyrus samples from reference and AD cases. Three CD36-positive endothelial populations were identified (Endo 1, Endo 2, and Endo 3). Endo 2, characterized by intermediate CD36 expression, was the predominant CD36-positive endothelial population in both reference and AD samples. Descriptively, AD samples showed a lower relative proportion of Endo 2 and a higher proportion of Endo 3 than reference samples (Figure 1A and Supplementary Figure 1A). Complementing the single-nucleus observations, we examined endothelial CD36 localization and arteriolar morphology in human AD brain tissue. In hippocampal and cortical specimens, CD36 immunoreactivity was prominent in arteriolar endothelium and was spatially heterogeneous along abnormal vessel walls. α-smooth muscle actin (α-SMA) staining showed irregular and discontinuous mural coverage with focal loss of circumferential smooth-muscle organization, while DAPI staining revealed disruption of the normally ordered vascular-wall nuclear pattern. Together, these observations document endothelial CD36 expression in morphologically abnormal arterioles in human AD tissue (Figure 1 and Supplementary Figure 1).

**Figure 1.**
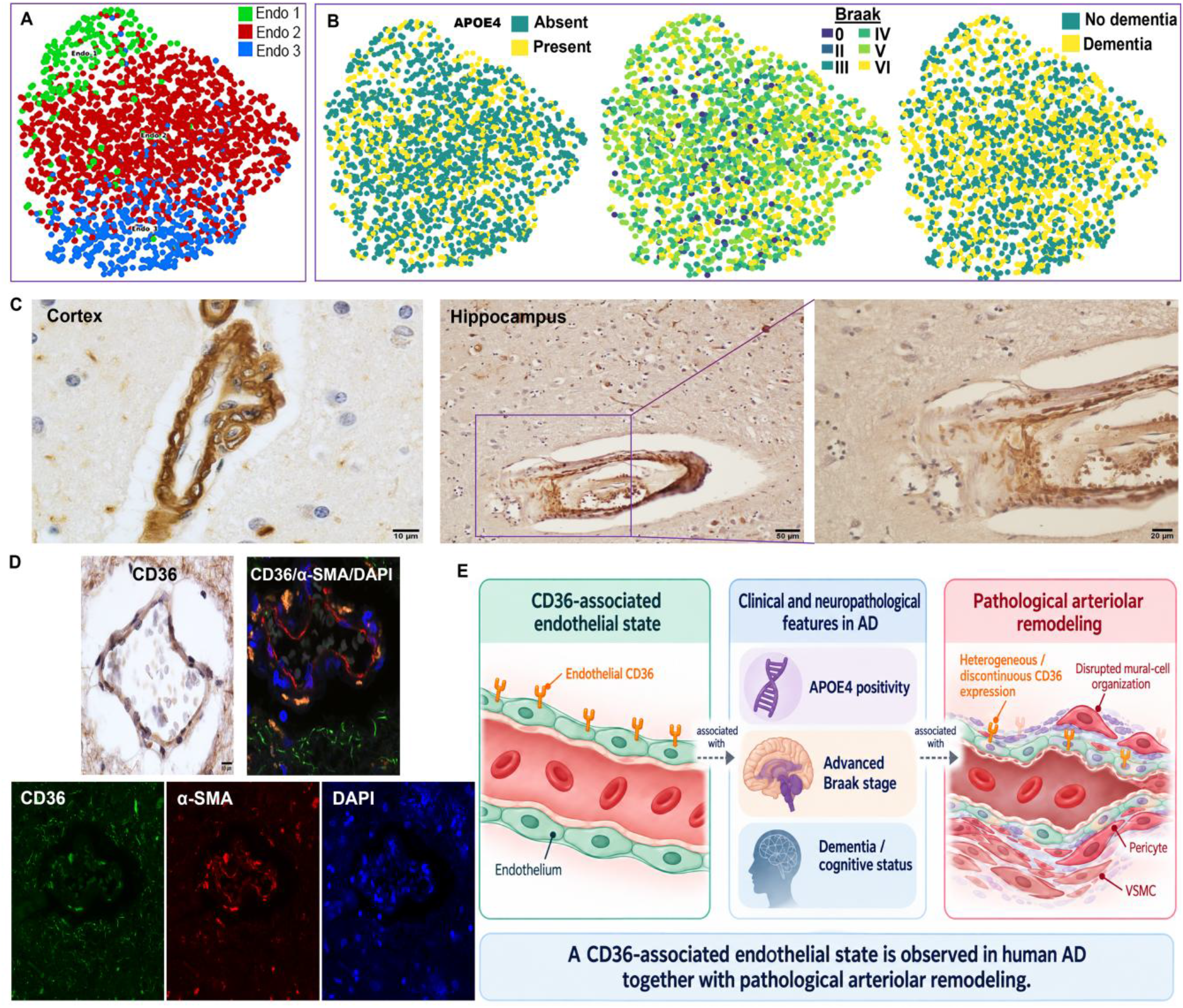
CD36-expressing endothelial heterogeneity and abnormal arteriolar morphology in human AD. **A.** UMAP visualization of CD36-expressing endothelial populations (Endo 1, Endo 2, and Endo 3) from the SEA-AD single-nucleus RNA-sequencing dataset. **B**. The same UMAP annotated with APOE4 status, Braak stage, and dementia status. **C**. Representative CD36 immunohistochemistry of AD cortex and hippocampus showing spatially heterogeneous staining in morphologically abnormal arterioles; the right panel enlarges the boxed hippocampal vessel. **D**. CD36 immunohistochemistry and CD36/α-SMA/DAPI immunofluorescence from AD brain tissue, showing heterogeneous vascular CD36 signal and disrupted α-SMA-positive mural organization. E. Conceptual illustration of possible relationships among CD36-associated endothelial states, AD-related clinical characteristics, and arteriolar pathology; these relationships are proposed for further investigation.

### Control and PSEN1 E276A iPSC-derived ECs show distinct transcriptional profiles and responses to GS4898

Having identified CD36-expressing endothelial heterogeneity and abnormal CD36-positive arterioles in human AD, we next asked whether endothelial transcriptional states could respond to pharmacologic perturbation. We differentiated control and PSEN1 E276A iPSC lines into ECs, confirmed endothelial identity using CD31 staining and endothelial-marker expression, and profiled vehicle-, 2 μM-, and 5 μM-GS4898-treated cells across three independent differentiation experiments. Global transcriptomic analysis separated control and PSEN1 E276A ECs, with cell-line background contributing substantially to variation along the first principal component. Both lines maintained endothelial identity but differed in baseline chemokine and vascular-remodeling transcriptional programs (Figure 2A and Supplementary Figure 2; Supplementary Table 1).

**Figure 2.**
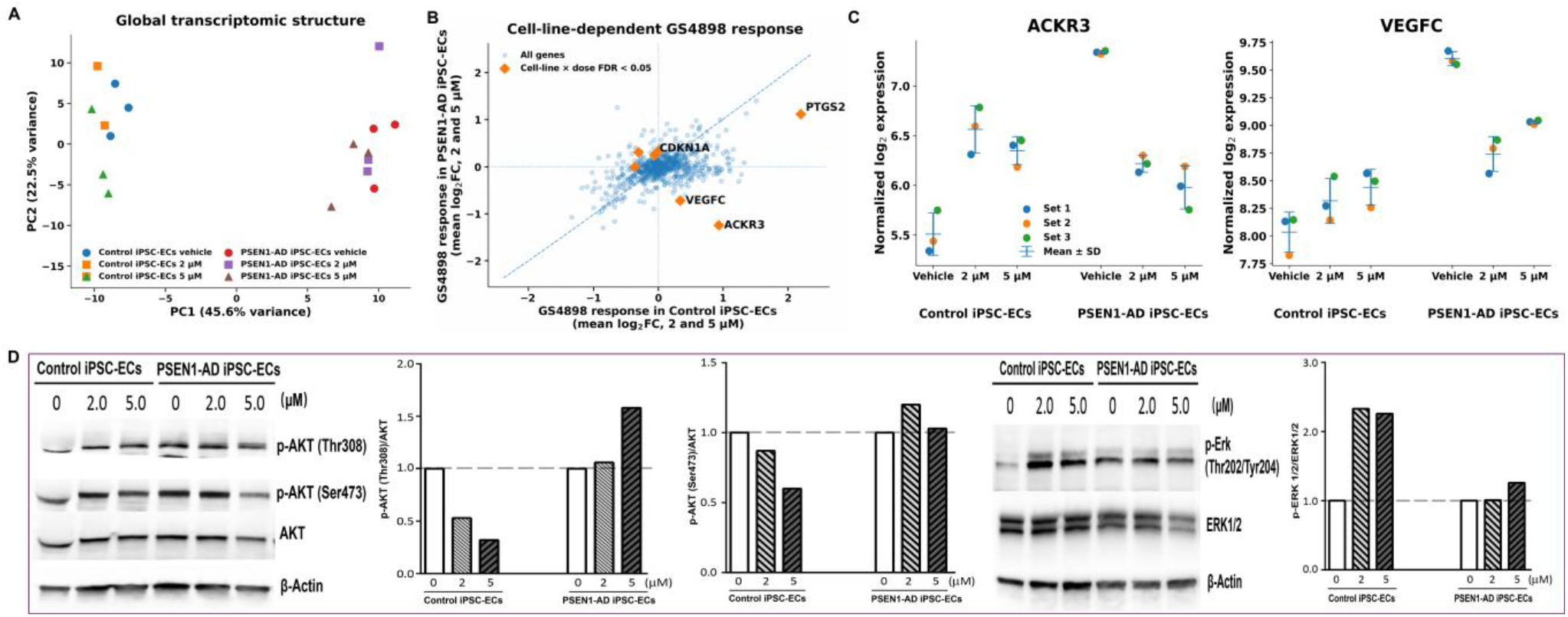
Control and PSEN1 E276A iPSC-derived ECs exhibit distinct baseline transcriptional profiles and responses to GS4898. **A.** Principal component analysis of NanoString profiles from control and PSEN1 E276A iPSC-ECs treated with vehicle, 2 μM GS4898, or 5 μM GS4898 across three independent differentiation experiments (18 samples total). **B**. Comparison of mean transcriptional responses to GS4898 (2 and 5 μM versus vehicle) in the two cell-line backgrounds. Each point represents a gene; orange diamonds identify significant cell-line-background-by-dose interactions following FDR correction. Representative genes include ACKR3, VEGFC, CDKN1A, and PTGS2. **C**. Normalized NanoString expression of ACKR3 and VEGFC across treatment groups. Points represent independent differentiation experiments; summary values are mean ± SD. **D**. Exploratory acute AKT and ERK signaling responses after 30 min of GS4898 treatment. Immunoblots show phospho-AKT (Thr308 and Ser473), total AKT, phospho-ERK1/2 (Thr202/Tyr204), total ERK1/2, and β-actin. Densitometric values are ratios of phospho-to total protein relative to vehicle. Repeated immunoblot runs using the same lysate were technical replicates, not independent biological replicates.

GS4898 elicited both shared and cell-line-dependent transcriptional responses. Whereas some programs changed similarly in control and PSEN1-AD ECs, ACKR3 and VEGFC showed divergent responses, illustrating how the two cellular backgrounds differed in their responses to the same perturbation (Figure 2B–C). Targeted RT-qPCR further supported this cell-line-dependent transcriptional response, revealing distinct patterns of endothelial-state marker regulation in control and PSEN1-AD iPSC-ECs (Supplementary Figure 3). The selected markers included SOX17, ACKR1, CXCL12, and HEY2. Exploratory RT-qPCR analysis suggested a trend toward reduced CD36 mRNA expression in control iPSC-ECs following treatment with 5 μM GS4898, although this change did not reach statistical significance (data not shown).

Exploratory acute signaling analysis also suggested distinct responses to GS4898. Control ECs showed increased ERK1/2 phosphorylation accompanied by reduced AKT phosphorylation, whereas PSEN1-AD ECs displayed a blunted ERK response and a different pattern of AKT regulation (Figure 2D). Together, these data indicate that PSEN1-AD endothelial cells remain transcriptionally responsive to GS4898, with a response distinct from that of control cells.

### Genetic models connect endothelial PKD1-CD36 regulation with cerebrovascular pathology and metabolic–immune programs

The iPSC-EC findings demonstrate pharmacologic transcriptional responsiveness and suggest a potential link between GS4898 treatment and CD36 expression. To further investigate the role of endothelial CD36 in cerebrovascular abnormalities and AD-relevant metabolic–immune regulation, we examined complementary genetic mouse models. Building on our previous findings that PKD1 signaling suppresses endothelial CD36 and promotes proarteriogenic reprogramming, 8,14,15 we examined cerebrovascular morphology and endothelial CD36 expression in endothelial-specific Pkd1-deficient mice. We observed that loss of endothelial PKD1 was accompanied by increased endothelial CD36 and marked cerebrovascular abnormalities, including abnormal vascular architecture and collagen deposition (Figure 3). These findings provided a vascular context for examining how CD36 itself regulates brain endothelial molecular programs.

**Figure 3.**
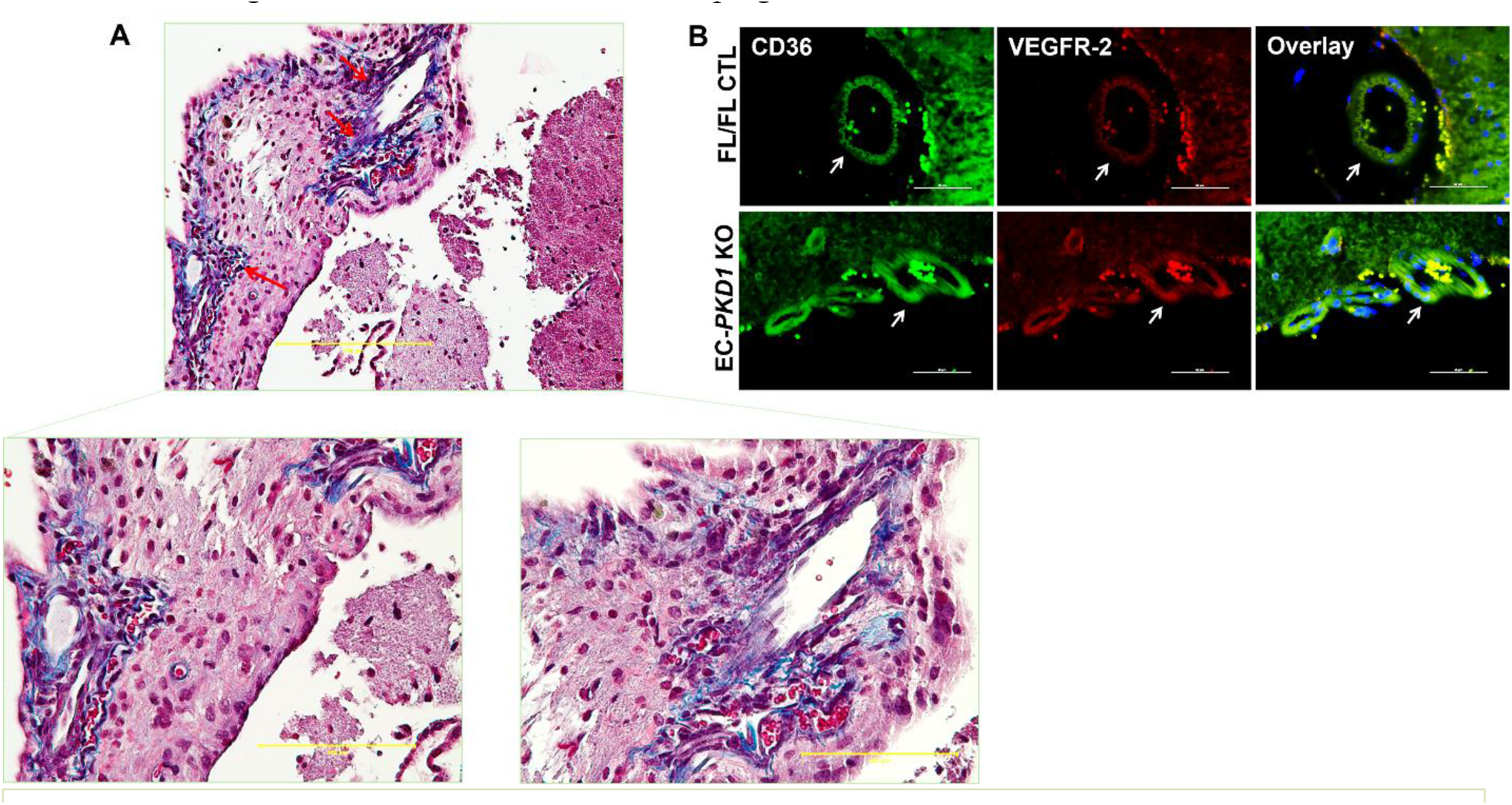
Endothelial PKD1 deficiency is associated with cerebrovascular abnormalities and increased endothelial CD36 expression. **A.** Representative trichrome staining of brain tissue from endothelial-specific Pkd1-deficient mice showing abnormal cerebrovascular architecture and collagen deposition. Lower panels are enlarged views of the indicated vascular regions. **B**. Representative immunofluorescence of CD36 and VEGFR-2 in control and endothelial-specific Pkd1-deficient brain vessels. Increased endothelial CD36 is observed in abnormal vessels from Pkd1-deficient mice. Arrows indicate selected vascular regions. Scale bars: 200 μm (upper trichrome), 100 μm (enlarged trichrome), and 50 μm (immunofluorescence).

To determine how endothelial CD36 itself regulates molecular programs relevant to brain vascular pathology, we analyzed brain endothelial ribosome-associated transcripts from endothelial-specific CD36-deficient TRAP mice exposed to a seven-week high-fat-diet metabolic-inflammatory challenge. This paradigm was used to reveal CD36-dependent endothelial responses under systemic metabolic stress rather than as a complete model of AD. RNA sequencing and pathway analysis showed broad transcriptional reorganization involving cytokine, interleukin, and inflammatory-response pathways (Figure 4). Functional clustering further identified immune-regulatory processes among differentially expressed genes, indicating that endothelial CD36 shapes inflammatory transcriptional programs in vivo.

**Figure 4.**
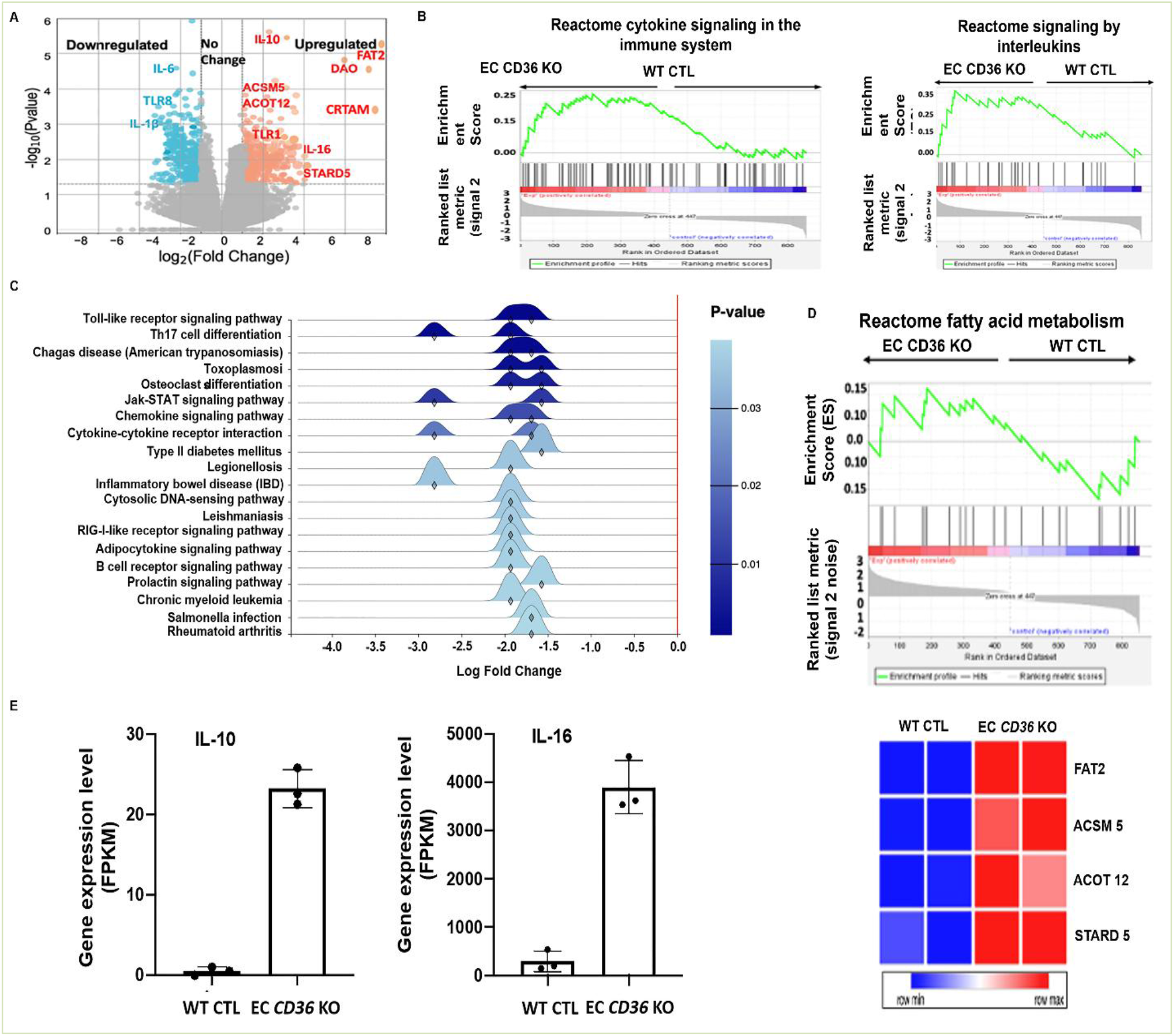
Endothelial CD36 deficiency alters immune- and metabolism-associated transcriptional programs in brain endothelium. Ribosome-associated transcripts from endothelial-specific CD36-deficient TRAP and control mice were analyzed following a seven-week high-fat-diet challenge. **A.** Volcano plot of differential gene expression with selected immune- and metabolism-associated genes. **B**. Gene set enrichment analyses of Reactome cytokine signaling and interleukin pathways. **C**. Pathway-level analysis of downregulated immune-memory-associated genes, including Toll-like receptor, Th17, JAK–STAT, chemokine, and cytokine signaling. **D**. Gene set enrichment of fatty-acid metabolic genes and heatmap of FAT2, ACSM5, ACOT12, and STARD5. **E**. RNA-seq expression of IL10 and IL16 in control and CD36-deficient brain endothelium; points represent samples, and bars show mean ± SD.

We further examined transcriptional programs associated with trained immunity, or innate immune memory. Analysis using the trained-immunity database identified multiple immune-memory-associated genes that were reduced in CD36-deficient brain endothelium, including marked downregulation of IL22. Network and pathway analyses connected CD36-regulated genes to Toll-like receptor, Th17, chemokine, and JAK–STAT signaling, with CCL3, IL22, and NFKBIA linking these pathways. Notably, disease-annotation analysis linked CD36-regulated endothelial genes to Alzheimer’s disease as well as systemic lupus erythematosus and broader immune-system pathways (Supplementary Figure 4D). Several AD-relevant genes, including HLA-DQA1, APOE, ACVRL1, C1QA, and APP, were reduced in CD36-deficient brain endothelium (Supplementary Figure 4E). These findings suggest broad remodeling of innate-immune-memory-associated transcriptional programs after endothelial CD36 loss rather than a change confined to one inflammatory pathway.

Importantly, these CD36-dependent, AD-annotated changes complement the human observations in Figure 1 and support a relationship between endothelial CD36 and molecular programs relevant to AD, without establishing disease specificity. Because CD36 is a major regulator of endothelial fatty-acid uptake and lipid metabolism, we next asked whether these immune changes were accompanied by changes in metabolic transcriptional programs.

Gene set enrichment analysis showed coordinated alteration of fatty-acid/lipid and amino-acid metabolic programs in CD36-deficient brain endothelium. Endothelial CD36 loss was accompanied by reduced IL1B, IL6, and TLR8 expression and increased IL10 and IL16. KEGG-based analyses further connected CD36-regulated fatty-acid metabolic genes with innate immune signaling pathways. Together with the AD-related gene and pathway annotations, these results show that endothelial CD36 loss reshapes lipid-metabolic and innate-immune-associated transcriptional programs with potential relevance to brain vascular pathology in AD (Figure 4 and Supplementary Figure 4).

## Discussion

Our study brings together three complementary lines of evidence for a CD36-centered endothelial-state framework in AD. First, human transcriptomic analyses identify heterogeneous CD36-expressing endothelial populations, while histopathology documents endothelial CD36 in structurally abnormal arterioles. Second, genetic mouse models place the PKD1-CD36 regulatory axis in a cerebrovascular context and show that endothelial CD36 regulates metabolic and innate-immune-associated transcriptional programs, including AD-relevant molecular signatures, under metabolic-inflammatory stress. Third, control and PSEN1 E276A iPSC-derived ECs remain responsive to GS4898, with distinct transcriptional responses across the two genetic backgrounds. These results connect vascular pathology, endothelial immunometabolic regulation, and pharmacologic responsiveness within a proposed endothelial-state framework, suggesting a potentially important role for endothelial CD36 in the vascular component of AD pathogenesis. However, they do not establish a single causal pathway from CD36 to AD progression or demonstrate drug-mediated vascular rescue.

The human transcriptomic and histopathological findings provide distinct yet complementary evidence relevant to the role of endothelial CD36 in AD-associated vascular pathology. SEA-AD analyses show that CD36 is expressed across several endothelial populations, with a lower relative proportion of Endo 2 and a higher proportion of Endo 3 among CD36-positive ECs in AD than in reference samples. Separately, tissue analysis identifies CD36 expression within morphologically abnormal arterioles. The SEA-AD Endo 2 population has not been spatially matched to these vessels; nevertheless, the observations together motivate investigation of CD36-associated vascular-state heterogeneity in AD. Although CD36 is commonly used as a capillary/metabolic endothelial marker in single-cell vascular atlases, its expression and phenotypic association are context dependent and may vary with endothelial state and tissue microenvironment.22,23 Prior cerebrovascular studies localized CD36 to cerebral ECs, showed increased endothelial CD36 in Tg2576 mice, and demonstrated CD36-dependent oxidative stress in penetrating arterioles and cerebral-amyloid-angiopathy-associated arteriolar dysfunction.9,10 Together with our human AD histology, these observations suggest that endothelial CD36 expression may vary with vascular segment, microenvironment, and disease-associated endothelial state. Such endothelial-state variation along the arteriovenous continuum is biologically plausible.4,24 Spatially resolved analyses may further clarify whether CD36-positive endothelial populations include transitional states and their vascular distribution in AD. 25

Importantly, CD36 may provide a mechanistic interface between endothelial metabolism and innate immune regulation. In addition to its established role in long-chain fatty-acid uptake, CD36 functions as a scavenger and signaling receptor capable of integrating extracellular lipid cues with cellular metabolic and inflammatory responses.5–7 Moreover, endothelial CD36 may function as a context-dependent regulator of vascular immunometabolic homeostasis. Under diet-induced metabolic-inflammatory stress, endothelial CD36 deletion reduced IL1B and IL6 expression while reshaping lipid-metabolic, innate-immune-memory-associated, and AD-relevant transcriptional programs. These findings suggest that CD36 coordinates endothelial metabolic and immune responses rather than acting solely as a proinflammatory receptor, consistent with the established relationship between immunometabolism and innate immune memory.13,26–28 Importantly, human studies have linked midlife obesity and vascular risk factors to subsequent AD risk and visceral obesity to inflammation-associated cerebral small-vessel abnormalities.29,30 Together with experimental evidence connecting obesity to immune dysregulation in AD and endothelial CD36 to cerebrovascular inflammation,31,32 our findings raise the possibility that excessive CD36 signaling promotes pathological inflammation and enhances endothelial immune memory.

Metabolic reprogramming is a core component of innate immune memory,^26^ and persistent innate immune adaptation involves coordinated metabolic and epigenetic changes.^13^,^27^,^28^ Recent work in intestinal barrier epithelia further demonstrates that nonhematopoietic barrier cells can shape tissue-resident immune memory through antigen presentation and sustained bidirectional interactions with immune cells.^33^ In this context, our findings suggest that CD36-dependent metabolic regulation may influence innate-immune-memory-associated endothelial programming through coordinated metabolic and immune transcriptional remodeling. Whether and how CD36 signaling contributes to maladaptive endothelial immune memory and neurovascular immune dysregulation in AD remain important questions for future investigation.

Beyond defining CD36-regulated programs, a distinct translational question is whether ECs with different genetic backgrounds retain pharmacologic responsiveness. Interestingly, control and PSEN1-AD iPSC-derived ECs maintained endothelial identity but occupied distinct transcriptional states, and GS4898 elicited both shared and cell-line-dependent responses, including divergent regulation of ACKR3 and VEGFC. These findings highlight endothelial transcriptional plasticity and suggest that pharmacologic modulation may depend on pre-existing endothelial states. Given the established role of AKT–ERK signaling balance in arterial morphogenesis and arteriogenesis,17,34 the differential signaling responses observed here may offer insights into the mechanisms underlying this plasticity. These observations support further exploration of state-selective therapeutic strategies targeting cerebrovascular dysfunction in AD.

Several limitations should be considered. The endothelial CD36-deficient TRAP studies were conducted under diet-induced metabolic-inflammatory stress rather than in an AD model. Although these genetic studies demonstrate CD36-dependent regulation of brain endothelial metabolic and immune transcriptional programs in vivo, their direct contribution to AD pathology remains to be established. The human histological analysis included a limited number of AD specimens without matched non-AD controls. Moreover, the SEA-AD and TRAP datasets do not resolve CD36-associated transcriptional programs specifically to the abnormal arterioles observed in human AD tissue or determine whether metabolic and immune alterations coexist within the same endothelial populations. The iPSC-EC studies used repeated differentiations of one control line and one CRISPR-engineered PSEN1 E276A line with distinct genetic backgrounds. Because these lines are not isogenic, the observed differences cannot be attributed specifically to the PSEN1 variant. The AKT–ERK signaling findings were exploratory, and the direct molecular target of GS4898 remains unknown. Finally, the present studies did not establish functional restoration of blood– brain barrier integrity, cerebral perfusion, arteriolar remodeling, or functional trained immunity. Nevertheless, the complementary human, genetic, and pharmacologic findings provide a coherent framework for investigating CD36-associated endothelial-state regulation and its therapeutic potential in AD.

## Author contributions

AK performed the endothelial TRAP transcriptomic and bioinformatic analyses, contributed to data interpretation, and participated in manuscript writing. JQZ performed the iPSC-EC NanoString, qPCR, and Western blot signaling studies, analyzed and interpreted the resulting data, and contributed to manuscript writing. JH performed human AD tissue histology and immunostaining, contributed to vascular image analysis and data interpretation, and participated in manuscript writing. RG contributed to selected data analysis and figure preparation. RC contributed pathology expertise and interpretation of human AD tissue findings. SP provided the iPSC lines and contributed to manuscript review and writing. AWB contributed vascular expertise and manuscript review. BR conceived the central scientific concept, directed the study, interpreted the data, and wrote and revised the manuscript. All authors reviewed and approved the manuscript.

## Acknowledgements

The work in the Laboratory of Molecular and Vascular Pathobiology has been supported by the Transformational Project Award (24TPA1293301) from the American Heart Association, National Institutes of Health (R01 HL136423), the Breast Cancer Research Foundation of Alabama, and the COVID-19 Pandemic Bridge Fund from the UAB Heersink School of Medicine (B.R.). RNA sequencing was performed by the UAB Heflin Center for Genomic Sciences. Human brain tissue samples and diagnostic data from patients with Alzheimer’s disease were provided by the UAB Tissue Biorepository and the Alzheimer’s Disease Research Center (P30AG086401). We thank Dr. Abdellah Akil for tissue endothelial RNA extraction and Dr. Jinjin Hao for establishing the high-fat-diet mouse model. We also thank Dr. Xiaofeng Yang for helpful discussions and suggestions regarding endothelial cell immunity, and Dr. Randall T. Peterson, University of Utah College of Pharmacy, for kindly providing GS4898.OpenAI’s ChatGPT provided substantial assistance with conceptual synthesis, integration of multidisciplinary evidence, literature analysis, manuscript restructuring and editing, and refinement of the endothelial-state framework. Microsoft Copilot and Google Gemini provided additional independent critical reviews of selected manuscript drafts.

## Supplementary figures and legend

**Supplementary Figure 1.**
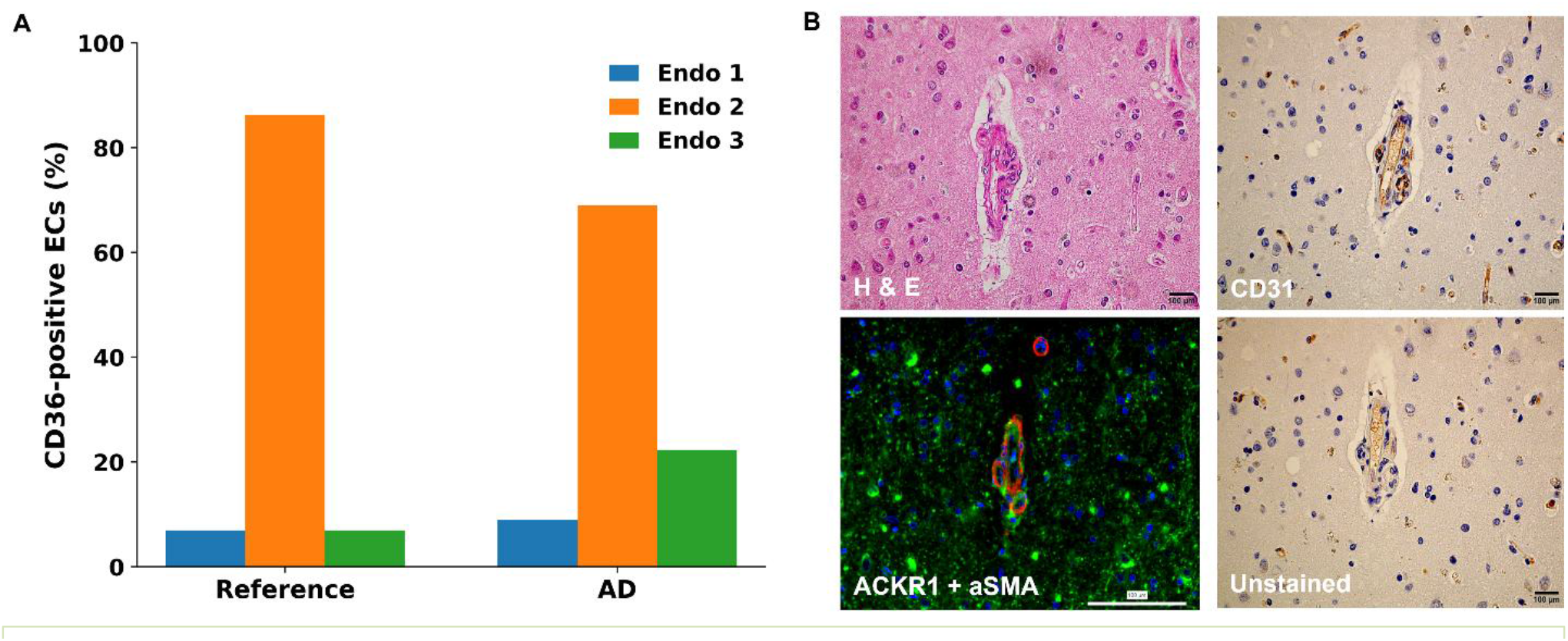
CD36-positive endothelial heterogeneity and vascular context in human AD. **A**. Relative distribution of CD36-positive endothelial populations in reference and AD middle temporal gyrus samples. Endo 2 constituted the predominant CD36-positive endothelial population in both groups. Values are shown as percentages of total CD36-positive endothelial cells within each group. **B**. Complementary histological and vascular characterization of human AD cortex. Representative H&E staining, CD31 immunohistochemistry, ACKR1/α-SMA immunofluorescence, and an unstained control provide vascular and histological context for the abnormal cortical vessels shown in Figure 1. Images were obtained from separate tissue sections and are not serially matched. Scale bars, 100 μm.

**Supplementary Figure 2.**
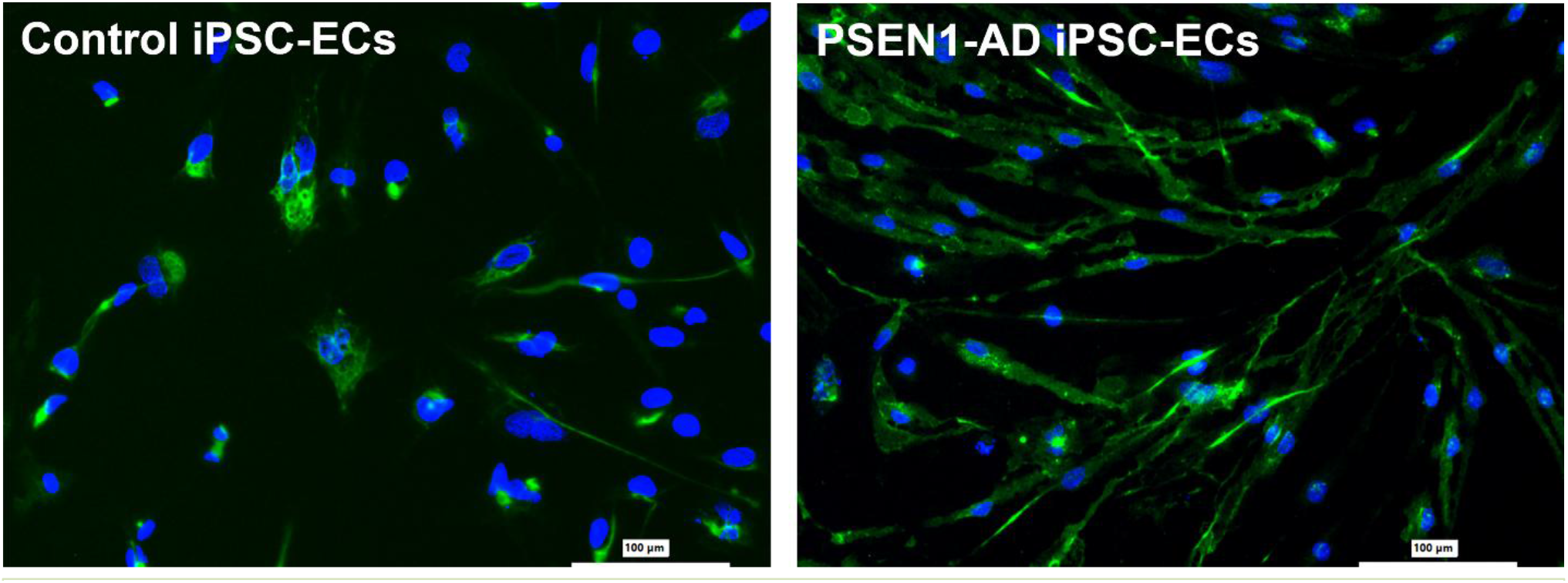
Endothelial identity of control and PSEN1-AD iPSC-derived endothelial cells. Representative CD31 immunofluorescence in control and PSEN1-AD iPSC-derived endothelial cells. CD31 is shown in green and nuclei are counterstained with DAPI (blue), supporting endothelial identity in both cell backgrounds. Scale bars, 100 μm.

**Supplementary Figure 3.**
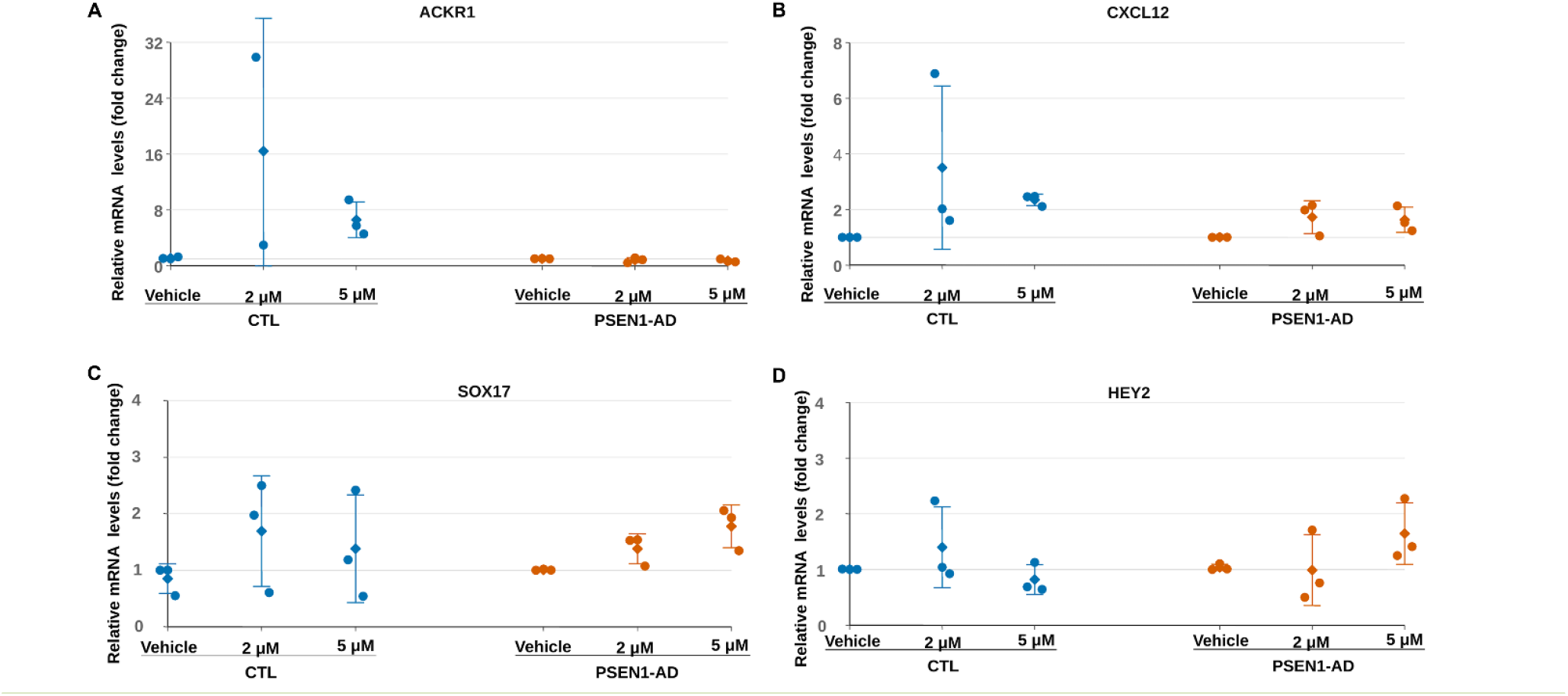
Targeted RT-qPCR analysis of GS4898-responsive endothelial-state markers in control and PSEN1-AD iPSC-derived endothelial cells. **A**. ACKR1. **B**. CXCL12. **C**. SOX17. **D**. HEY2. Relative mRNA expression was normalized to GAPDH and expressed relative to the matched vehicle condition. Technical triplicates were averaged within each differentiation experiment before analysis. Individual circles represent independent differentiation sets; diamonds and error bars indicate mean ± SD. Three differentiation sets were available for each condition except ACKR1 in control cells treated with 2 μM GS4898, for which n = 2.

**Supplementary Figure 4.**
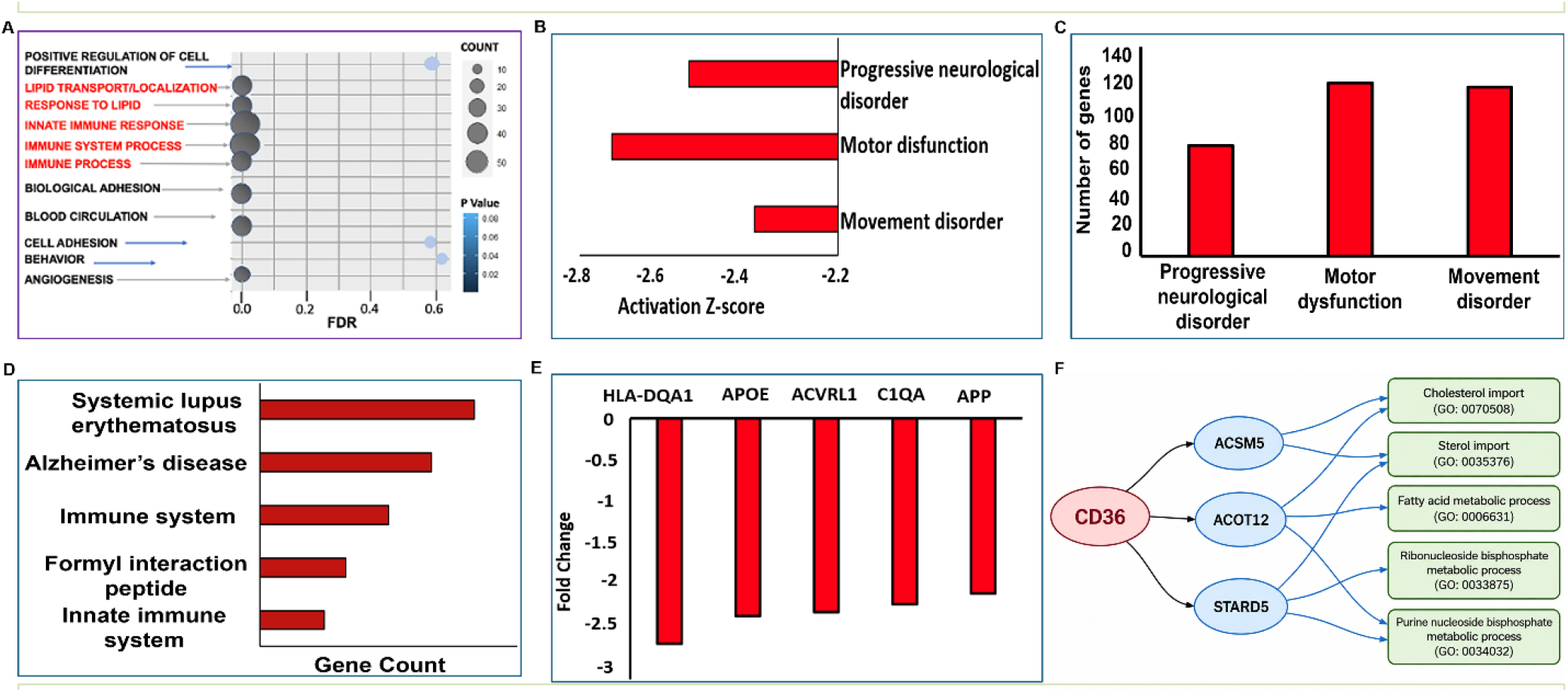
Extended functional, disease-associated, and metabolic analyses of the CD36-dependent brain endothelial transcriptome. **A**. Functional enrichment analysis of CD36-regulated endothelial genes showing biological processes related to lipid transport and response, innate and immune responses, blood circulation, cell adhesion, and angiogenesis. **B**. Functional annotation of differentially expressed genes showing activation Z-scores for progressive neurological disorder, motor dysfunction, and movement disorder. **C**. Numbers of differentially expressed genes associated with these neurological-disorder categories. **D**. Gene-count analysis showing associations of CD36-regulated genes with systemic lupus erythematosus, Alzheimer’s disease, immune-system, innate-immune-system, and related annotation categories. **E**. Expression changes in selected neurological- and Alzheimer’s disease-associated genes, including HLA-DQA1, APOE, ACVRL1, C1QA, and APP. **F**. Functional network analysis further characterizing metabolic reprogramming associated with endothelial CD36 deficiency. Consistent with the fatty-acid metabolic changes shown in Figure 4D, network analysis generated using Enrichr and refined using Cytoscape and NDEx links CD36-regulated fatty-acid metabolism-associated genes, including ACSM5, ACOT12, and STARD5, to cholesterol import, sterol import, fatty-acid metabolism, and related metabolic processes.

**Supplementary Table 1.**
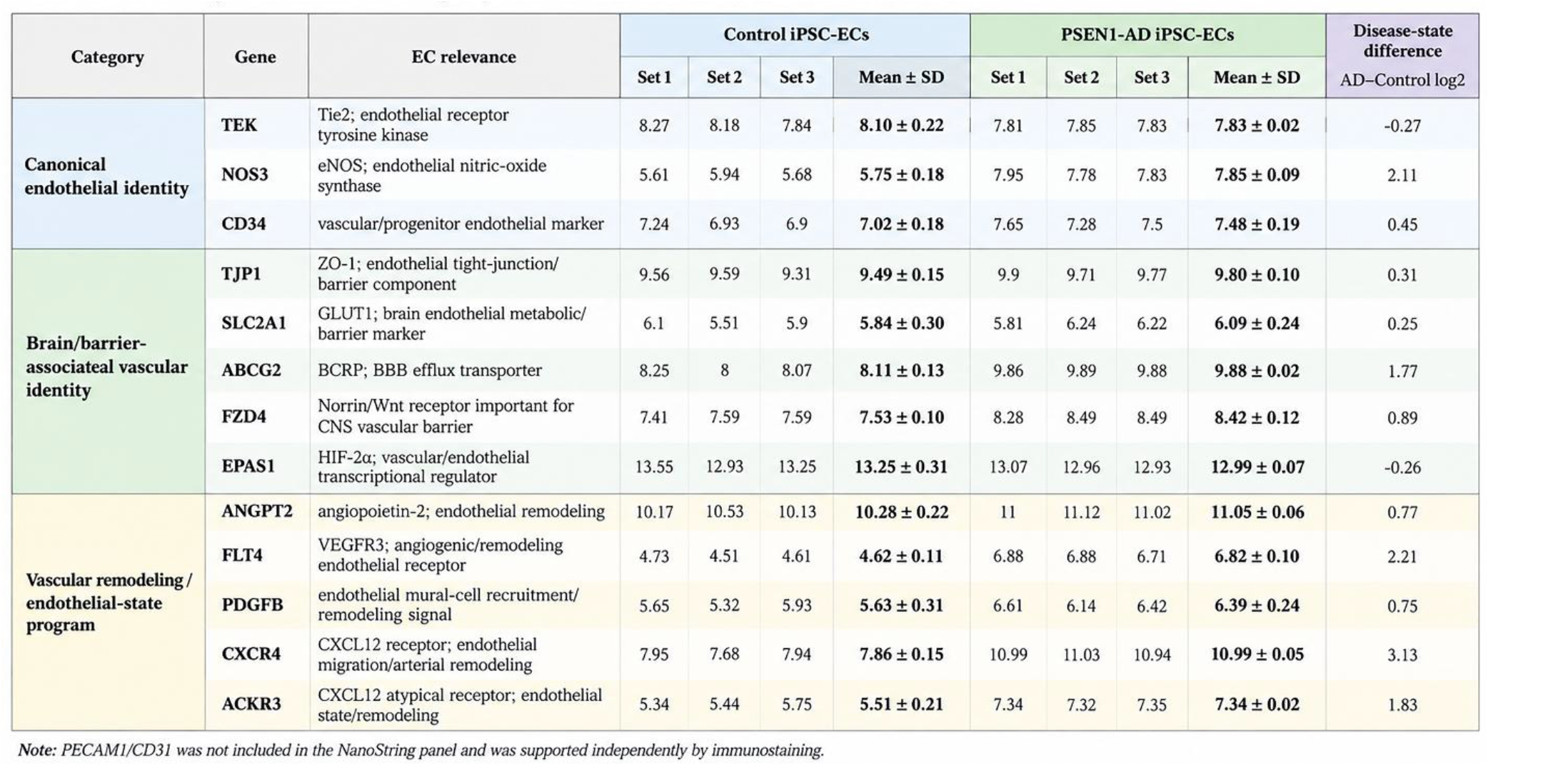
NanoString endothelial-marker expression in vehicle-treated iPSC-derived ECs. Baseline NanoString endothelial-marker expression in control and PSEN1-AD iPSC-derived ECs. Baseline samples without GS4898 treatment were analyzed. Values are normalized log2 expression. Control = A0/C0/E0; PSEN1-AD = B0/D0/F0. Markers are grouped as canonical endothelial identity, brain/barrier-associated vascular identity, or vascular-remodeling/endothelial-state programs. PECAM1/CD31 was not included in the NanoString panel; endothelial identity was supported independently by CD31 immunostaining (Supplementary Figure 2).

